# Artificial intelligence to solve the X-ray crystallography phase problem: a case study report

**DOI:** 10.1101/2021.12.14.472726

**Authors:** Irène Barbarin-Bocahu, Marc Graille

## Abstract

The determination of three dimensional structures of macromolecules is one of the actual challenge in biology with the ultimate objective of understanding their function. So far, X-ray crystallography is the most popular method to solve structure, but this technique relies on the generation of diffracting crystals. Once a correct data set has been obtained, the calculation of electron density maps requires to solve the so-called «phase problem » using different approaches. The most frequently used technique is molecular replacement, which relies on the availability of the structure of a protein sharing strong structural similarity with the studied protein. Its success rate is directly correlated with the quality of the models used for the molecular replacement trials. The availability of models as accurate as possible is then definitely critical.

Very recently, a breakthrough step has been made in the field of protein structure prediction thanks to the use of machine learning approaches as implemented in the AlphaFold or RoseTTAFold structure prediction programs. Here, we describe how these recent improvements helped us to solve the crystal structure of a protein involved in the nonsense-mediated mRNA decay pathway (NMD), an mRNA quality control pathway dedicated to the elimination of eukaryotic mRNAs harboring premature stop codons.

## Introduction

The central dogma of molecular biology implies the transfer of the information contained within genes into the corresponding proteins. Although the amino acid sequence of a protein is important, the three dimensional structure is even more crucial for a protein to fulfill its cellular and biochemical functions. Indeed, misfolding or aggregation due to point mutations or other causes are known to be responsible for many pathologies including neurodegenerative disorders such as Alzheimer’s disease (Dobson, 2003, Forman et al., 2004). Hence, since the resolution of the three dimensional structure of myoglobin (Kendrew et al., 1958), the first one to be determined, extensive efforts have been devoted to solve the structures of proteins from various organisms. As a consequence, almost 180,000 structures are currently deposited at the Protein Data Bank, a 50-years old database (Berman et al., 2000). The ambitious structural genomics initiatives launched in the early 2000’s after the release of the first complete genome sequences, have also participated to this effort. In addition to the determination of at least 15,000 new structures, these programs have contributed to the development of high-throughput strategies for the cloning, expression, purification, crystallization and structure determination of proteins of interest, that are now implemented in many structural biology departments (Terwilliger et al., 2009, Grabowski et al., 2016, Michalska & Joachimiak, 2021). The PDB is then a fantastic catalogue of structures that can be used for homology-based protein structure prediction by molecular modeling. This is one of the holy grail in biology so as to help researchers and clinicians to appreciate protein biological and biochemical functions as well as the potential impact of mutations associated with diseases. Consequently, extensive efforts have been devoted for many decades to the development of protein structure prediction approaches either by template-based or template-free modeling. Improvements were constantly realized but very recently, a breakthrough step has been made as testified by the results of the recent Critical Assessment of protein Structure Prediction (CASP14) experiment (Pearce & Zhang, 2021). Indeed, the accuracy of the models obtained by the machine learning method AlphaFold developed by the DeepMind company was impressive compared to those obtained by other methods (Jumper et al., 2021, Lupas et al., 2021, Pereira et al., 2021). This inspired the improvement of the RoseTTaFold program by the Baker’s lab (Baek et al., 2021). The greatly improved accuracy of the structure predictions generated by AlphaFold and RoseTTAFold then offers the promise that many of these *in silico* three dimensional models can be good to excellent templates for researchers to conduct further studies.

These more accurate models also open great opportunities to accelerate the structure determination process by X-ray crystallography. Indeed, during the crystal diffraction experiment, the intensities of the individual diffracted X-ray waves are recorded but the information related to their phases is lost. Hence, one major hurdle encountered by structural biologists is to obtain these phases through various approaches such as multiple isomorphous replacement (MIR) using heavy atoms derivatives, single or multiple wavelength anomalous diffusion (SAD or MAD) mostly from crystals of selenomethionine-substituted proteins, molecular replacement and in some specific cases *ab initio* phasing (Rupp, 2009). The molecular replacement method is by far the most popular as around 70% of the crystal structures currently deposited in the PDB were determined using this technique. It relies on the use as search model of the structure of a protein sharing strong structural similarity (*i*.*e*. in general more than 30% sequence identity) with the crystallized protein (Abergel, 2013, Scapin, 2013). Hence, its success rate is directly correlated with the quality of the template used. One can then expect that the more accurate models predicted by either AlphaFold or RoseTTAFold will strongly facilitate and accelerate the determination of protein structures by the molecular replacement technique.

Here, we describe how the recent improvements in structure modeling helped us to solve the crystal structure of the *Kluyveromyces lactis* orthologue of the Nmd4 protein (hereafter named *Kl*Nmd4). In *Saccharomyces cerevisiae* yeast, Nmd4 is involved in the nonsense-mediated mRNA decay pathway (NMD), an mRNA quality control pathway dedicated to the elimination of eukaryotic mRNAs harboring premature stop codons (He & Jacobson, 1995, Dehecq et al., 2018). The *Kl*Nmd4 protein is predicted to be made of a single PIN (for PilT N-terminus) domain, which is endowed with RNA binding activity and in many cases harbors endonuclease activity (Senissar et al., 2017). Although we obtained crystals of the *Kl*Nmd4 protein diffracting up to 2.45Å resolution, we struggled for almost 2 years trying to solve its structure. Traditional approaches were tried unsuccessfully. We soaked crystals with heavy metals solutions but obtained no derivative. As *Kl*Nmd4 only contains the initiator methionine, we introduced additional methionines by site directed mutagenesis for SAD/MAD phasing using selenomethionine but none of these mutants crystallized. We also performed molecular replacement trials using various models but none of them gave clear solutions. Based on the above mentioned recent advances in protein structure prediction, we have used both AlphaFold and RoseTTAFold programs to generate new structural models for the *Kl*Nmd4 protein. We show that the quality of these models lead to rapid determination of the structure of *Kl*Nmd4 while models generated using few other programs do not.

## Materials and methods

### Cloning, protein over-expression and purification

To enhance the expression yield of *Kluyveromyces lactis* Nmd4 (*Kl*Nmd4; UniProt ID Q6CVZ8), we expressed it as a fusion protein with a N-terminal His_6_-ZZ double tag (where ZZ stands for two Z domains from the IgG binding *Staphylococcus aureus* protein A (Nilsson et al., 1987)). We first inserted a *de novo* synthesized DNA sequence (Integrated DNA Technologies, Belgium) encoding for the His_6_-ZZ tag followed by a 3C protease cleavage site into the pET28-b plasmid using *Nco*I and *BamH*I restriction enzymes to generate the pET28-His_6_-ZZ-3C plasmid. Next, the DNA sequence encoding the *Kl*Nmd4 protein was amplified by polymerase chain reaction using the genomic DNA of the NK40 strain (generous gift from Dr K. Breunig) as a template together with oligonucleotides oMG593 and oMG594 (Table S1). This PCR product was cloned into the pET28-His_6_-ZZ-3C plasmid using *BamH*I and *Xho*I restriction enzymes to generate plasmid pMG897 (Table S1).

The *Kl*Nmd4 protein was expressed in *Escherischia coli* BL21(DE3) Gold strain (Agilent technologies) using pMG897 plasmid and 1 L of auto-inducible terrific broth media (ForMedium AIMTB0260) containing kanamycin (50 μg/mL) first for 4 hours at 37°C and then overnight at 25°C. Cells were harvested by centrifugation (4,000 g at 4°C for 45 min) and resuspended in buffer A (20 mM Tris-HCl pH 7.5, 5 mM β-mercaptoethanol, 200 mM NaCl). The cells were lysed by sonication on ice in the presence of 200 μM phenylmethylsulfonyl chloride (PMSF) protease inhibitor and the lysate cleared by centrifugation at 20,000 g at 4°C for 45 min. The supernatant was loaded onto Ni–NTA Sepharose High Performance affinity resin (GE Healthcare Biosciences) pre-equilibrated with buffer A. The resin was then washed extensively with buffer A followed by two more washing steps with buffer A supplemented respectively with 1 M NaCl first, and then 20 mM imidazole pH 7. The recombinant His_6_-ZZ tagged *Kl*Nmd4 protein was eluted with buffer A supplemented with 400 mM imidazole pH 7. The His_6_-ZZ tag was cleaved overnight under dialysis conditions (buffer B : 20 mM Hepes pH 7, 200 mM NaCl, 5 mM β-mercaptoethanol) upon addition of 3C protease (70 μL at 4 mg/mL). The protein was then loaded onto HiTrap SP Fast Flow column (GE Healthcare Biosciences) and eluted with a linear gradient of buffer B from 50 mM NaCl to 1M NaCl. The fractions containing *Kl*Nmd4 were then applied onto a S75-16/60 size-exclusion chromatography column (GE Healthcare Biosciences) using buffer B (GE Healthcare Biosciences) and a flow rate of 1 mL/min. The fractions containing pure *Kl*Nmd4 were collected and concentrated to 15 mg/mL.

### Size exclusion chromatography-multi-angle laser light scattering (SEC-MALLS)

The *Kl*Nmd4 protein (100 μL at 1 mg/mL) was injected at a flow rate of 0.75 mL/min on a Superdex^™^ 200 Increase 10/300 GL column (GE-Healthcare) using buffer B. Elution was followed by a UV-visible spectrophotometer, a MiniDawn TREOS detector (Wyatt Technology) and a RID-20A refractive index detector (Shimadzu). Data were processed with the program ASTRA 6.1 (Wyatt Technology). The *M*w was directly calculated from the absolute light scattering measurements using a dn/dc value of 0.183.

### Crystallization, data collection and processing

Crystallization trials were performed by mixing 150 nL of protein with an equal volume of different crystallization solutions in 96-well TTP Labtech plates at 7°C using the Mosquito automate (TTP Labtech). Prior to X-ray exposure, the crystals were transferred into the crystallization solution supplemented with 20% ethylene glycol and 20% glycerol and flash-cooled in liquid-nitrogen. The data were collected at 100 K on Proxima-2a beamline (Synchrotron SOLEIL, Saint-Aubin, France). Several datasets collected from a single crystal were processed using the XDS program, merged and scaled using the XSCALE program (Kabsch, 1993).

## Results

### Purification and crystallization of Kluyveromyces lactis Nmd4

Initially, we tried to crystallize the Nmd4 protein from *Saccharomyces cerevisiae* but did not obtain crystals. We then decided to focus on the Nmd4 protein from the *Kluyveromyces lactis* yeast (*Kl*Nmd4), which shares 38.6% sequence identity and 55.2% sequence homology with the Nmd4 protein from budding yeast. *Kl*Nmd4 has been purified using a three steps purification procedure as described in the materials and methods section. The purified *Kl*Nmd4 protein was analyzed by SEC-MALLS, revealing that it exists as a monomer in solution (measured molecular weight of 28.0 kDa *versus* theoretical molecular weight of 28.2 kDa; Fig. 1A). We next obtained rhombohedral crystals in the following crystallization condition : 0.1 M sodium citrate pH 5.6, 0.9-1 M Li_2_SO_4_ and 0.6 M ammonium sulfate, within one day at 7°C. These crystals appeared within 6 days and grew for at least two more weeks to reach 200 μm length (Fig. 1B). They diffracted on the Proxima-2a beamline from the French synchrotron SOLEIL in 2019 (Duran et al., 2013). Taking advantage of their size and of this microfocus beamline, we collected several datasets from the same crystal. By merging three datasets together, we obtained a 2.45 Å resolution dataset (Table 1). These crystals belong to the P321 space group or its related enantiomorphs (P3_1_21 or P3_2_21) with an estimated number of two *Kl*Nmd4 molecules per asymmetric unit assuming a 50% solvent content (Matthews, 1968).

**Table 1:**
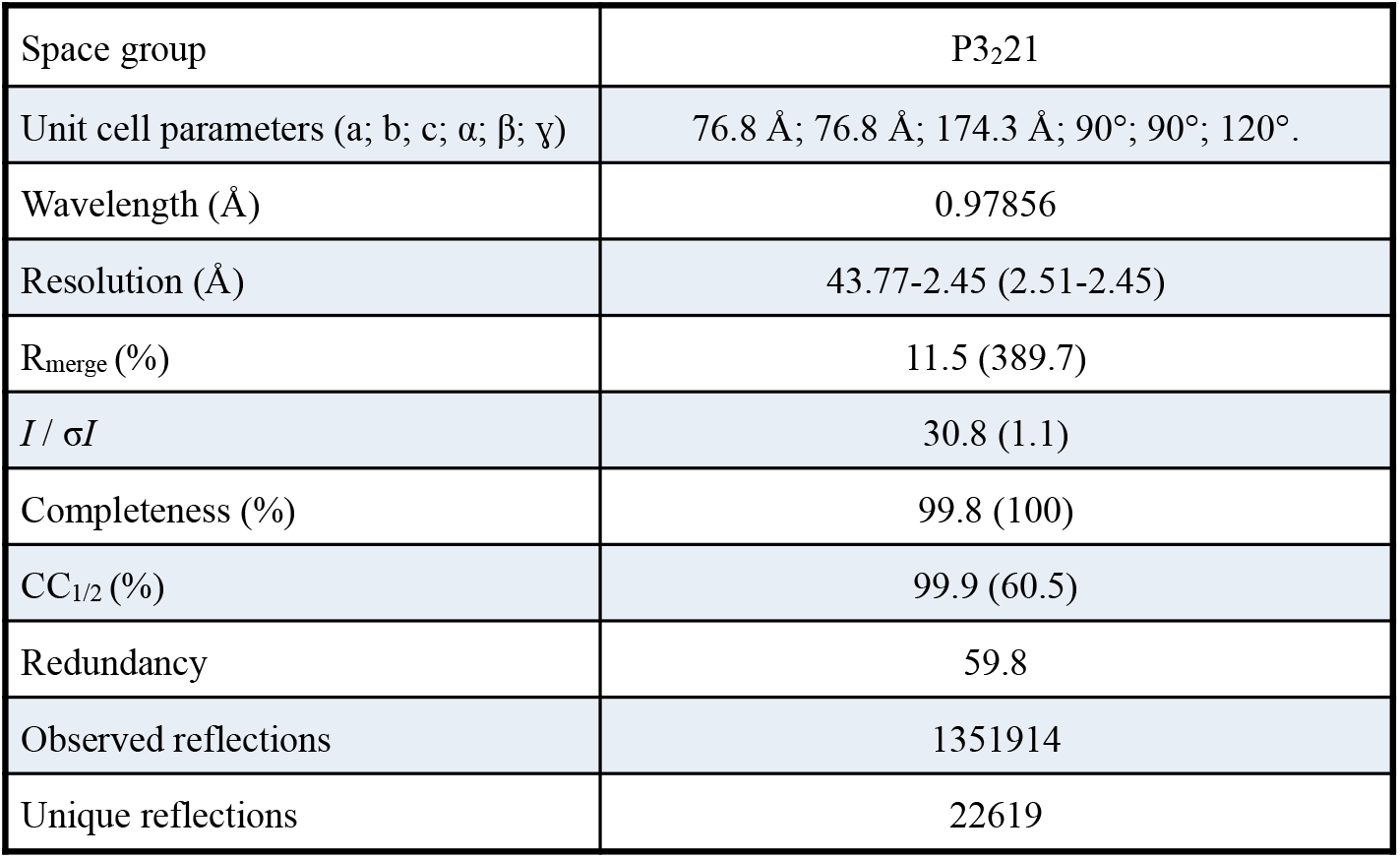
Data collection statistics.

**Figure 1 :**
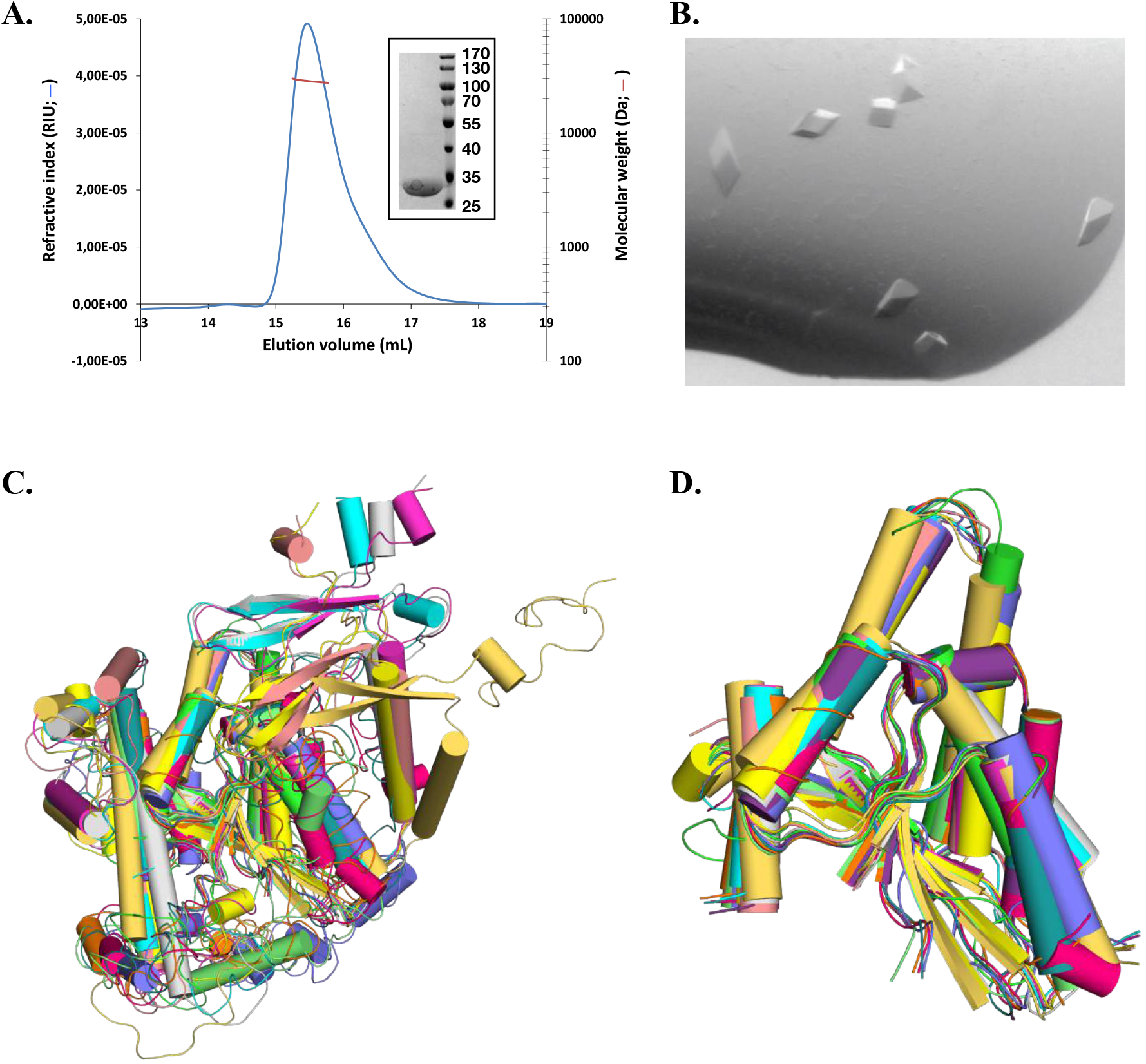
*Kl*Nmd4 characterization and crystals. A. The *Kl*Nmd4 protein is monomeric in solution. Zoom-in representation of the only peak visible on the SEC-MALLS chromatogram obtained from *Kl*Nmd4. The refractive index is shown as a blue line (left y-axis) while the distribution of molecular mass calculated from light scattering along this peak is shown in red (right logarithmic y-axis). Inset : SDS-PAGE analysis of the Coomassie-stained proteins used for this experiment. The molecular weight (kDa) of the ladders is indicated on the right of the inset. B. Rhombohedral crystals of the *Kl*Nmd4 protein. C. Superimposition of all the intact models generated for the *Kl*Nmd4 protein. The *α-*helices and β-strands are shown as cylinders and arrows, respectively. Each model is shown with a different color. D. Superimposition of all the *Kl*Nmd4 models obtained after trimming of the most divergent regions and used for molecular replacement assays. Same color code as in panel C.

### Generation of KlNmd4 structure models

As no protein of known three-dimensional structures shared more than 25-30% sequence identity and as *Kl*Nmd4 contains no methionine, we tried to solve the structure of this protein by multiple isomorphous replacement following soaking of the crystals into the crystallization solution supplemented with different heavy atom salt solutions. We also tried to introduce extra methionine residues by mutating hydrophobic amino acids but these variants were clearly less stable than the native protein and consequently, none of them crystallized. In parallel, we performed molecular replacement assays using several models for the *Kl*Nmd4 protein generated with a set of programs or servers available two years ago : PHYRE-2 (Kelley et al., 2015), Swiss-Model (Waterhouse et al., 2018), RosettaCM (Song et al., 2013) and i-Tasser (Yang et al., 2015). For PHYRE-2 model, we selected the one generated using the crystal structure of the PIN domain from human SMG6 (PDB code 2HWW; (Glavan et al., 2006)) as template (predicted sequence identity with *Kl*Nmd4 of 22%). The Swiss-Model server generated a single model from another crystal structure of the same protein domain (PDB code 2DOK; (Takeshita et al., 2007)). The i-Tasser server used both structures to generate five different models while we selected the *ab initio* mode for the RosettaCM server to also generate five different models (when several models have been generated using the same software, there are annotated from a to e). More recently, we generated one and five additional models using the AlphaFold and RoseTTAFold softwares, respectively (Baek et al., 2021, Jumper et al., 2021).

In all these 18 *in silico* models, the *Kl*Nmd4 protein is predicted to be made of a single PIN domain. The superimposition of all these models reveals that the structural core made of *α-*helices and β-strands is well conserved but that the conformations of most of the loops connecting secondary structure elements strongly vary from one model to another one (Fig. 1C). We then removed these loops to generate truncated search models, to avoid the rejection of correct solution during the molecular replacement trials due to a too high number of inter-molecular steric clashes caused by these loops in the crystal packing (Fig. 1D). This strategy is commonly used for molecular replacement. With the exception of the Swiss-Model (residues 1-2 and 176 to 185 absent) and PHYRE-2 (residues 1-5 and 176 to 185 lacking) models, all the final models contain residues 1-80, 115-126 and 150-185. This grossly corresponds to the residues with per-residues confidence scores (pLDDT) higher than 90 in the AlphaFold model and to about half of the total amino acids of this protein.

### Structure determination by molecular replacement

For molecular replacement, we tried to position two copies of these different truncated models using two programs in parallel : MOLREP (version 11.7.02; (Vagin & Teplyakov, 1997)) and PHASER (version 2.8.3; (McCoy et al., 2007)). For each program, the default parameters as defined in the CCP4 interface (version 7.0.078; (Winn et al., 2011)) were used. For instance, MOLREP used the data included within the 43.76-3.02 Å resolution range while PHASER included all the data (*i*.*e*. up to 2.45 Å). In addition, we arbitrarily selected a rms difference of 0.75 Å between the models and the *Kl*Nmd4 crystal structure for PHASER. Here, we will discuss only the results obtained in space group P3_2_21 as it turned out to be the correct one. We tried each model individually, *i*.*e*. 18 different trials with each program. Finally, all the solutions obtained were refined using the BUSTER program (version 2.10.4; (Bricogne et al., 2017)) using 5 macrocycles of refinement and one TLS (for Translation-Libration-Screw) group per monomer.

#### MOLREP

To analyze the results for the different molecular replacement trials performed with MOLREP, we mostly focused on the contrast value calculated by the program as suggested in the tutorial. Basically, this value reflects the difference between the highest score and the mean score of the different solutions obtained after the translation fonction search has completed and the higher this score the more likely the solution is to be correct. In general, contrast values higher than 3 are strong indication that the proposed solution is correct. The distribution of the contrast values proposed by the MOLREP program for the different solutions obtained from the various models, clearly shows that the AlphaFold model largely outcompetes the other models and successfully led to the correct positioning of both *Kl*Nmd4 monomers (Fig. 2A). Indeed, the contrast values for the first (13) and second (9.5) copies of the AlphaFold model positioned are very high, indicating correct positioning. Next, the RoseTTAFold models yield some solutions with contrast scores higher that the threshold of 3. The theta, phi and chi angles of the rotation functions of these solutions are similar to those of the AlphaFold solutions, meaning that they are correctly positioned too. It is noteworthy that MOLREP could correctly position two copies for the RoseTTAFold-e and RoseTTAFold-a models while only one copy of the other RoseTTAFold models could be correctly positioned. The contrast values obtained with all the other models are mostly below 2, suggesting that these solutions are incorrect. Interestingly, deeper analysis of the results obtained with the other models indicates that for PHYRE-2 and i-Tasser-e models, one monomer was correctly positioned while the second one was not (Fig. 2A). However, the scores for these solutions were low and similar to the scores of incorrectly positioned monomers. This later observation clearly shows that although some monomers can be correctly positioned, it is difficult to identify them among all the wrongly positioned models. Finally, for those models where one molecule was correctly positioned, if we use 100 rotation functions in the translation function, *i*.*e*. more than when we use the default parameters defined in the MOLREP program, the correct position of the second monomer could only be found for the RoseTTAFold-b and RoseTTAFold-c models.

**Figure 2 :**
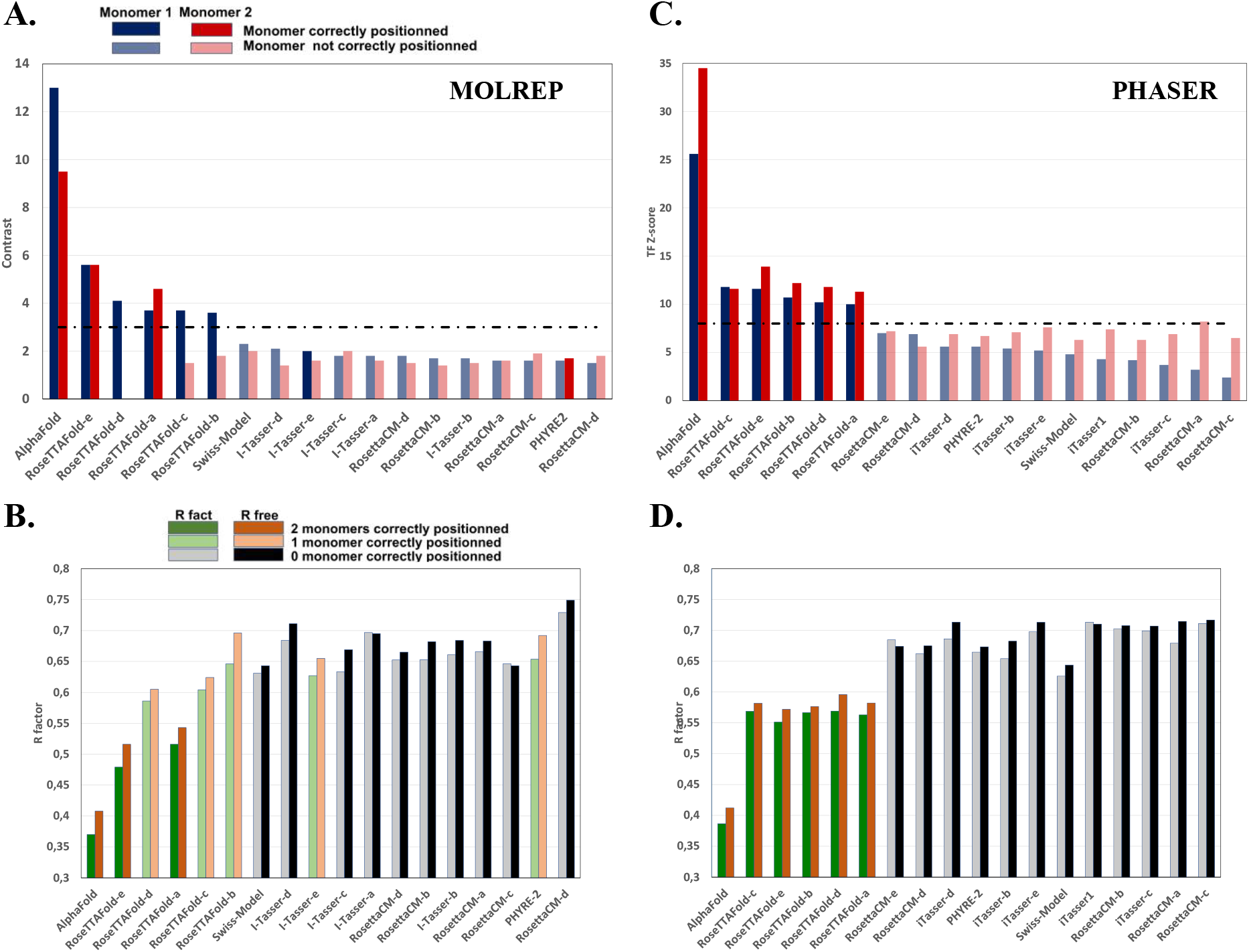
Comparison of the molecular replacement and refinement statistics obtained with different search models. A. Contrast values of the molecular replacement solutions obtained using the MOLREP program. Correct solutions are highlighted as dark colors while wrong solutions are in light colors. The color code is shown above the graph and was also used for panel C. The dashed line depicts the contrast value threshold above which solutions are considered to be correct. B. R and R_free_ values obtained after the refinement of the MOLREP solution obtained for each model. The color code is shown above the graph and is also valid for panel D. C. TF Z-scores of the molecular replacement solutions obtained using the PHASER program. The dashed line depicts the TF Z-score value above which solutions are considered to be correct. D. R and R_free_ values obtained after the refinement of the PHASER solution obtained for each model.

Next, we refined the coordinates of the solutions proposed for these different molecular replacement trials using the BUSTER program and analyzed the R and R_free_ values obtained for all these solutions (Fig. 2B). Once again, the solution obtained with the AlphaFold model outcompetes the other ones as testified by the much lower R and R_free_ values (37% and 40.8%, respectively) compared to the solutions obtained with RoseTTAFold-e (R and R_free_ values of 47.9% and 51.7%, respectively) and RoseTTAFold-a models (R and R_free_ values of 51.7% and 54.3%, respectively). As expected, the other solutions, *i*.*e*. with one or two monomers incorrectly positioned, yielded significantly higher R and R_free_ values (Fig. 2B), typical to those obtained for incorrect solutions.

#### PHASER

In parallel, we performed molecular replacement trials using the PHASER program and each of these 18 models. In that case, we analyzed the results by monitoring the TF Z-scores obtained for each solution. Indeed, it is considered that for the first positioned molecule, TF Z-score higher than 8 are indicative of correct solutions while TF Z-scores in between 6 and 8 indicate probable correct solutions (Phaserwiki). In our case, the solutions obtained using the AlphaFold model as well as the five models from RoseTTAFold yielded solutions with TF Z-scores higher than 10 for the first monomer and similar or higher for the second monomer (Fig. 2C). This indicates that for each of these 6 models, two copies were correctly positioned. On the contrary, the statistics of the solutions obtained with any of the other models suggested incorrect solutions and this was confirmed by the theta, phi and chi angles of the selected rotation functions, which significantly differ from those of the correct solutions.

The refinement of these different solutions with the BUSTER program revealed important gaps in the R and R_free_ values between these different solutions (Fig. 2D), which can then be divided into 3 groups. The first one with mean R and R_free_ values of 68.1% and 69.6%, respectively, corresponds to the incorrectly positioned molecules. The second group with mean R and R_free_ values of 56.4% and 58.2%, respectively, contains the solutions obtained with any of the RoseTTAFold models. Finally, the solution obtained using the AlphaFold model yielded R and R_free_ values of 38.7% and 41.2%, respectively, *i*.*e*. much better than those of any other solutions. The important gap between the R and R_free_ values calculated for the AlphaFold solution and for the RoseTTAFold solutions indicates that the RoseTTAFold models are probably more distant from the crystal structure than the AlphaFold model. It also suggests that the refinement/building cycles necessary to reach the final experimental structure will probably be more complex when starting from the RoseTTAFold models than from the AlphaFold model.

In summary, while the quality of the models generated two years ago using different protein structure prediction tools was insufficient to solve the crystal structure of *Kl*Nmd4, the models obtained with the recently implemented machine learning tools (AlphaFold and RoseTTAFold) were of much better quality and rapidly led to correct solutions using the two most popular molecular programs (PHASER and MOLREP). PHASER proved to be more efficient with the RoseTTAFold models as it correctly found two solutions for all the five tested models while MOLREP could only position correctly two copies for two out of the five RoseTTAFold models. Yet, the AlphaFold model yielded solutions with much higher scores than those obtained with RoseTTAFold models. Similarly, the statistics of the refinement of the solutions found using the AlphaFold model were much better than those of the RoseTTAFold models. Altogether, this suggests that the AlphaFold model is more similar to our experimental crystal structure than the RoseTTAFold models (see later).

### Structure of the KlNmd4 protein

Using the molecular replacement solutions obtained by MOLREP with the AlphaFold model, we performed iterative cycles of building and refinement at 2.45 Å resolution to converge to the final structure of the *Kl*Nmd4 protein (R and R_free_ values of 23.4% and 28.2%, respectively; Table 2). The quality of the 2Fo-Fc electron density map allowed us to model residues 1 to 128, 146-193 and 219-230 from monomer A and residues 1 to 130, 146-185 and 241-244 from monomer B. In addition, we could model a short pentapeptide from monomer A but it was not possible to assign it to a specific sequence of *Kl*Nmd4. Both monomers are virtually identical (rmsd of 0.241 Å over 145 Cα atoms) and hence, only the structure of monomer A will be discussed here.

**Table 2 :** Structure refinement statistics.

|  |  |
| --- | --- |
| Resolution (Å) | 43.77-2.45 |
| No. reflections | 22619 |
| R / R<sub>free</sub> (%) | 23.3 / 28.4 |
| *Number of atoms* |  |
| Protein | 2969 |
| Small molecules | 23 |
| Water | 7 |
| *B-factors (Å<sup>2</sup>)* |  |
| Wilson plot | 89.1 |
| Mean value | 95.4 |
| *R.m.s deviations* |  |
| Bond lengths (Å) | 0.009 |
| Bond angles (°) | 1.02 |
| PDB code | 7QHY |

The *Kl*Nmd4 protein is made of a central five stranded parallel β-sheet surrounded by 11 *α-*helices (Fig. 3A). Searches for proteins with high structural similarities identified human SMG6 PIN domain as closest hit (Fig. 3B; Z score of 12.4; rmsd of 2.5 Å over 160 Cα atoms and 20% sequence identity) as well as the PIN domains from several other archaeal and eukaryotic proteins (human SMG5, the RRP45 exosome subunit …), validating the bio-informatics analyses that led most protein structure prediction tools to use the human SMG6 structure as a template to model the *Kl*Nmd4 structure. The PIN domain from metazoan SMG6 proteins has been shown to be endowed with endonucleolytic activity (Glavan et al., 2006, Huntzinger et al., 2008, Eberle et al., 2009). This activity relies on the presence of three highly conserved acidic residues (D1251, D1353 and D1392 in human SMG6) at the heart of the active site (Fig. 3B). Importantly, these three acidic residues are not strictly conserved within fungal Nmd4 orthologues (Fig. 3C). This is particularly the case for D1353 from human SMG6, which has been shown to be critical for its endonucleolytic activity (Glavan et al., 2006, Eberle et al., 2009). The corresponding residue in Nmd4 proteins is hydrophobic (F114 in *Kl*Nmd4) as in human SMG5, a protein also devoid of endonuclease activity (Glavan et al., 2006). Altogether, this strongly argues in favor of the loss of this enzymatic activity in the fungal Nmd4 proteins (Fig. 3B-C).

**Figure 3 :**
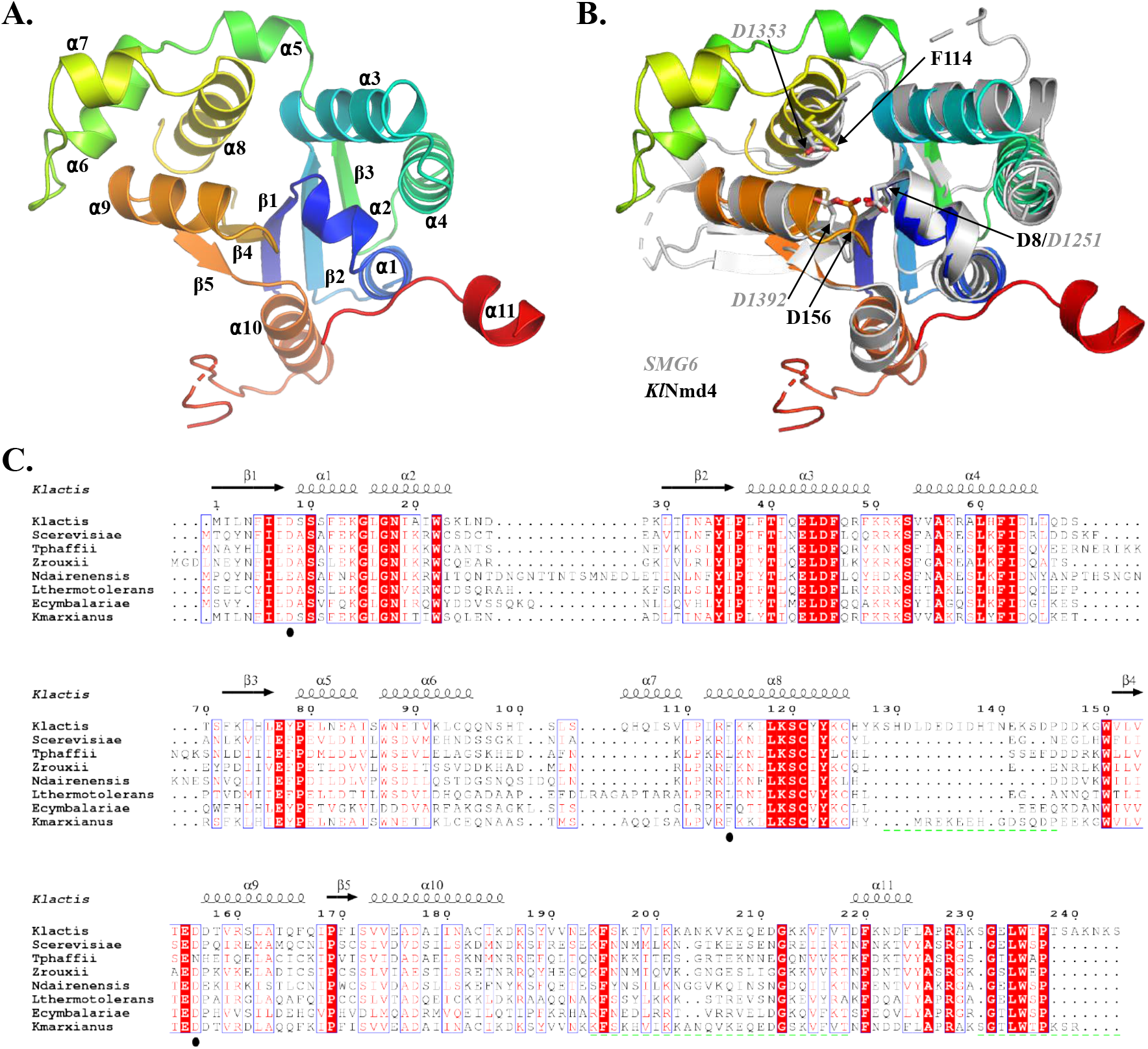
Structure of the *Kl*Nmd4 protein. A. Ribbon representation of the *Kl*Nmd4 crystal structure colored from its N-terminal (blue) to its C-terminal (red) extremities. B. Superimposition of human SMG6 PIN domain (grey) onto the *Kl*Nmd4 structure. The side chain of the human SMG6 residues forming the endonuclease active site are shown as grey sticks and the corresponding residues in *Kl*Nmd4 structure are also shown as sticks. Labels referring to SMG6 are shown in italics. C. Multiple sequence alignment of fungal Nmd4 proteins. Strictly conserved residues are in white on a red background. Partially conserved amino acids are in red and boxed. Undefined residues in the final *Kl*Nmd4 model are indicated by dashed green lines below the alignment. Secondary structure elements as observed in the crystal structures of the *Kl*Nmd4 protein are shown above the alignment. Position corresponding to the human SMG6 endonucleolytic active site are indicated by black circles below the alignement.

## Discussion and conclusion

The way from a purified protein to the determination of its three dimensional crystal structure is paved with two major hurdles, *i*.*e*. the obtention of crystals diffracting beyond 3-4 Å resolution and the solution of the phase problem. Here, we report the case of a protein crystal structure, which could be solved thanks to the tremendous progress recently made in the field of protein structure prediction by the AlphaFold and RoseTTAFold programs. Indeed, all our extensive efforts to solve this structure by multiple isomorphous replacement, anomalous diffusion and molecular replacement failed during almost 2 years. The AlphaFold and RoseTTAFold models played a major role as they gave correct molecular replacement solutions. In particular, the overall quality of the AlphaFold model led to a correct solution very rapidly. This solution exhibits much higher contrast scores with MOLREP and TF Z-scores with PHASER than those obtained with any other model as well as much lower R and R_free_ values after the direct refinement of the molecular replacement solution (Fig. 2). This is due to the overall excellent quality of the *Kl*Nmd4 model proposed by AlphaFold, which is definitely much more similar to the experimental crystal structure than any of the other generated models (Fig. 4A). Indeed, the rmsd value between the structural core of the PIN domain from the AlphaFold model (*i*.*e*. the model lacking the flexible loops and that mostly corresponds to regions with pLDDT scores higher than 90) and the final crystal structure is 0.43 Å over 127 Cα atoms (Fig. 4A-B). By comparison, the second model (RoseTTAFold-e) yielding to the best statistics (*i*.*e*. higher contrast and TF Z-score values and lower R and R_free_) exhibits a rmsd value of 1.4 Å over 121 Cα atoms. This higher rmsd value is mostly due to the slightly different position of many *α-*helices relative to the central β-sheet in the RoseTTAFold-e model compared to the final structure (Fig. 4C). In addition to this similarity observed for the structural core of the PIN domain, the comparison of the full-length AlphaFold model with the final crystal structure (rmsd of 0.97 Å over 165 Cα atoms) reveals that the prediction of the region encompassing residues 81-114 (corresponding to helices α5 to α7, which was initially removed from the search model) in the AlphaFold model is very similar to the conformation trapped in our crystal structure (Fig. 4B). The structure of this loop was predicted with relatively high confidence (70 < pLDDT < 90) by AlphaFold. The loop connecting helix α8 to strand β4 (residues 127 to 150), which was modeled with relatively low confidence (pLDDT scores lower than 50 for most residues of this loop), is not visible in our crystal structure, indicating intrinsic flexibility (Fig. 4B). Finally, the C-terminal region starting from lysine 185 adopts different conformation between the AlphaFold model and our crystal structure. In the AlphaFold model, this region is predicted to fold as a long *α-*helix followed by a two stranded β-sheet (Fig. 4B) but with an overall lower confidence score (pLDDT scores lower than 70 for most residues) than for the structural core of the PIN domain. In our structure, only part of this region could be modeled due to overall intrinsic flexibility and the modeled residues do not match with the AlphaFold model. We observe the same overall trend when comparing the RoseTTAFold-e model to the crystal structure (rmsd value of 1.65 Å over 151 Cα atoms) albeit with larger deviations than for the AlphaFold model (Fig. 4A).

**Figure 4 :**
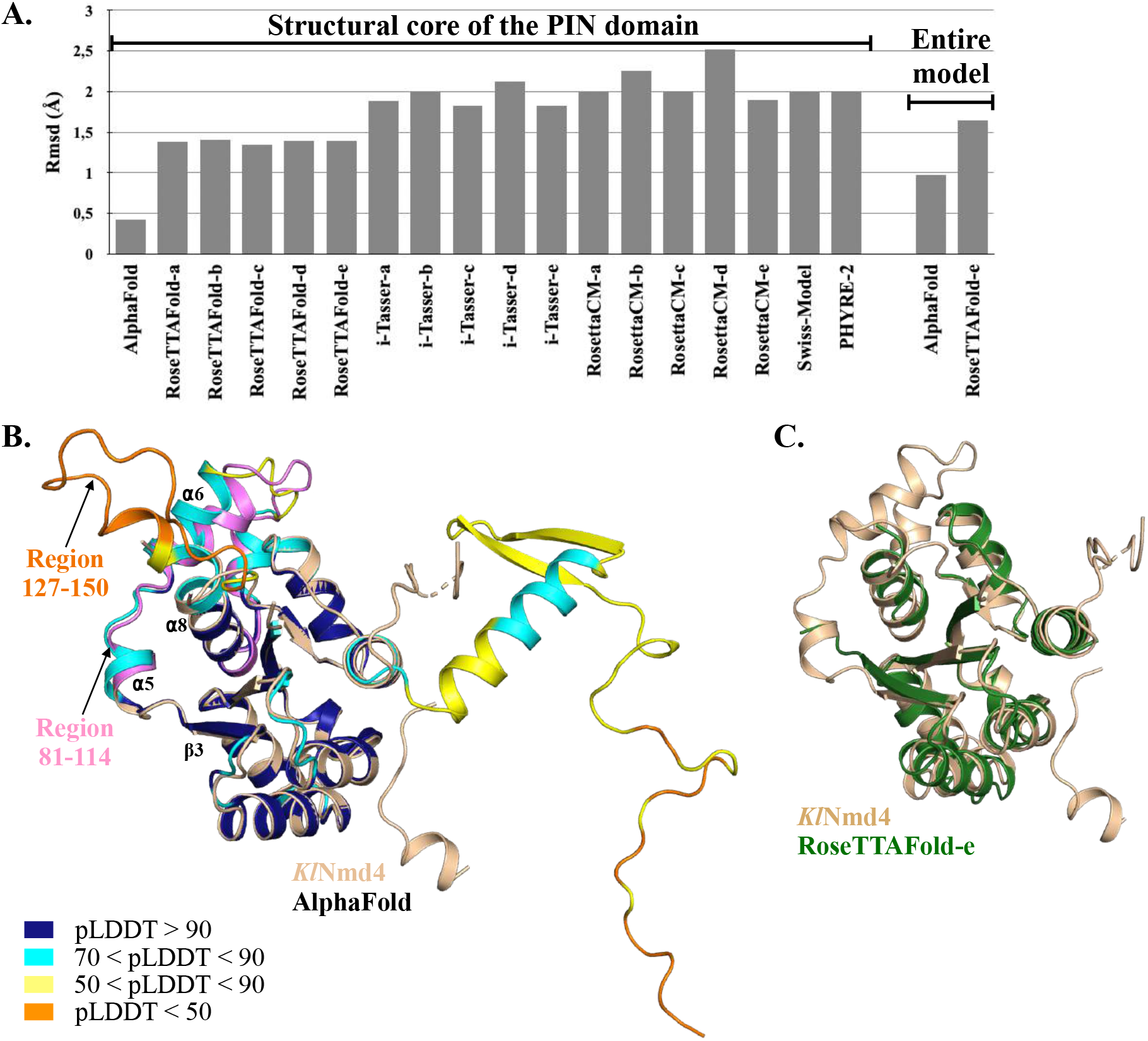
Comparison of *Kl*Nmd4 crystal structure with the different models. A. Graph depicting the rmsd value between the Cα atoms of the *Kl*Nmd4 crystal structure and of the different models either truncated (identified as «structural core of the PIN domain) or intact («entire model »). B. Superimposition of the full-length *Kl*Nmd4 AlphaFold model onto the *Kl*Nmd4 crystal structure (beige). This latter is colored according to the pLDDT values calculated by AlphaFold. The region 81-114 from the *Kl*Nmd4 crystal structure is highlighted in pink. C. Superimposition of the *Kl*Nmd4 crystal structure (beige) and the truncated *Kl*Nmd4 RoseTTAFold model (model e, dark green).

Interestingly, the *Kl*Nmd4 models obtained from RoseTTAFold can lead to correct structure solution using either MOLREP and PHASER, while none of the five models generated by its former version, RosettaCM yielded correct molecular replacement solution (Fig. 2). This is mostly due to the lower rmsd value of the structural core of the PIN domain of the various RoseTTAFold models (mean rmsd value of 1.38 Å) compared to the RosettaCM models (mean rmsd value of 2.18 Å; Fig. 4A), demonstrating a significant improvement in protein structure prediction with RoseTTAFold compared to RosettaCM. It is also noteworthy that for one i-Tasser model (i-Tasser-e) and for the PHYRE-2 model, MOLREP correctly positioned one monomer while PHASER did not (Fig. 2A and 2C). Unfortunately, the contrast scores calculated for these two solutions were comparable to those of incorrectly positioned solutions and hence, they did not emerge as correct ones. This is most likely due to the high rmsd values (1.8-2 Å) between both models and the final structure (Fig. 4A). This also illustrates the strong impact of model accuracy on the identification of correct molecular replacement solutions

Our results confirm the great improvements recently made in the prediction of protein structure field by the machine learning methods implemented in AlphaFold but also RoseTTAFold and suggest that the overall quality of these models will help in cracking the phase problem in many cases. This is in line with the outstanding results obtained by AlphaFold during the CASP14 session (Millan et al., 2021, Pearce & Zhang, 2021, Pereira et al., 2021). We managed to solve the structure of a small protein and to obtain 2Fo-Fc electron density maps of excellent quality within less than an hour while we were struggling for almost two years since the obtention of good quality diffraction datasets on this project. From discussions with other colleagues, we know that several other structural biologists have also succeeded in solving reluctant crystal structures thanks to the AlphaFold model of their favorite proteins. Other examples describing these successes will undoubtedly be published in the near future. Our simple routine can be easily automated as done using different molecular replacement and refinement programs in the CaspR server or MrBUMP program for instance (Claude et al., 2004, Keegan & Winn, 2008). As many other structural biology colleagues (Cramer, 2021, Perrakis & Sixma, 2021, Thornton et al., 2021), we are convinced that this achievement will definitely revolutionize structural biology. It is then very likely that the future protein and multi-protein complexes crystal structures will most largely be determined by molecular replacement and that methods such as MIR/SIR using heavy metal derivatives, SAD/MAD using selenomethionine-labelled proteins will become more and more marginal or applied to specific cases such as low resolution data.

This incredible breakthrough in the accuracy of predicted three dimensional structures opens amazing perspectives for biology in general. Indeed, thanks to the combined action of DeepMind and EMBL-EBI, every biologist has now access to the AlphaFold Protein Structure Database (https://alphafold.ebi.ac.uk/) presenting the models of the almost complete proteomes from 21 prokaryotic and eukaryotic model organisms including the *Methanocaldococcus jannaschii* archeon, some bacteria (*Escherichia coli, Mycobacterium tuberculosis…), Saccharomyces cerevisiae, Caenorhabditis elegans* and of course *Homo sapiens* (Tunyasuvunakool et al., 2021, Varadi et al., 2021). No doubt that this ressource is already fueling experiments in many biology labs. Very importantly, this database gives easy access to the confidence score (pLDDT) for each amino acid in the predicted structure. In our case, there is a strong agreement between these pLDDT values and the similarity between the AlphaFold model and our experimental structure (Fig. 4B). Among the regions from the *Kl*Nmd4 AlphaFold model with pLDDT values lower than 90, some adopt the same conformation as in the crystal structure, while others do not. These differences can be due to errors in the AlphaFold prediction, to the crystal packing, which can select a specific conformation, or to inherent flexibility. Hence, this pLDDT value reflects the confidence in the prediction but also the intrinsic flexibility of some protein regions. It is then important to consider these new *in silico* high quality models but also the crystal structures with some distance as they both contain biais (error in prediction, crystal packing effect…). It is then crucial to keep in mind that these three dimensional models need to be experimentally validated and should be questioned if the experimental data disagree with the model.

## Acknowledgements

We are grateful to the SOLEIL Synchrotron (France) staff (proposal numbers 20181001 and 20201046), in particular Martin Savko and Serena Sirigu (Proxima-2a beamline) for smoothly running the facility. MG acknowledges financial supports from the Centre National pour la Recherche Scientifique (CNRS), the Agence Nationale pour la Recherche (ANR; ANR-18-CE11-0003-04) and Ecole Polytechnique. IBB is supported by a PhD fellowship from the French Ministère de l’Enseignement Supérieur et de la Recherche (MESR).

## Author contributions

I.B.B. performed the experiments. I.B.B and M.G. designed research, analyzed the experiments and wrote the paper.

---

**Table S1.**
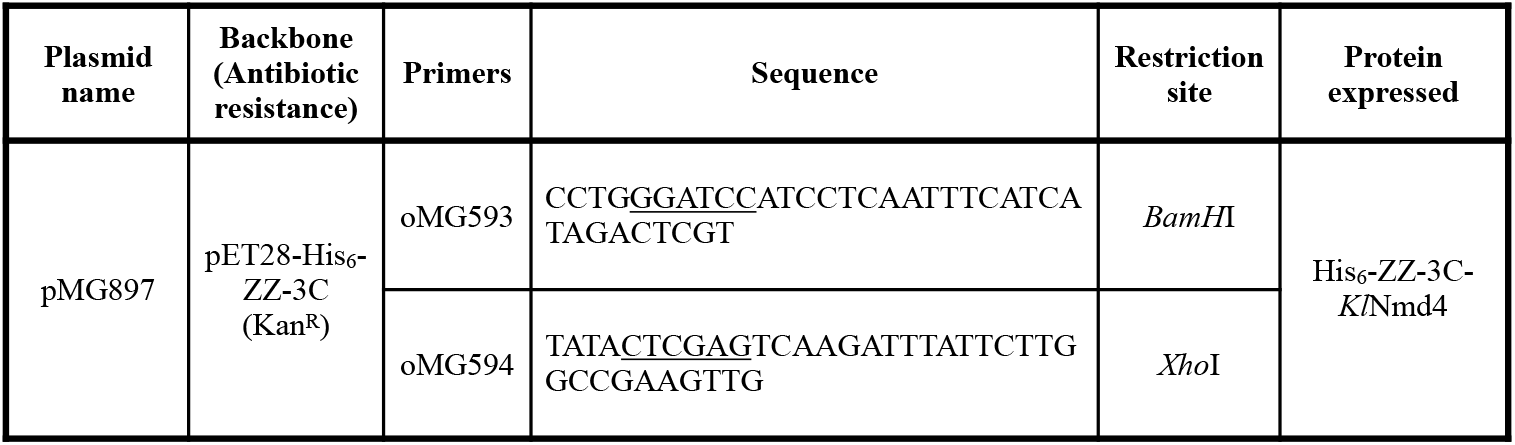
Primers and plasmid used for heterologous expression of *Kl*Nmd4.

